# Long-term Immunity of a Microneedle Array Patch of SARS-CoV-2 S1 Protein Subunit Vaccine Irradiated by Gamma Rays in Mice

**DOI:** 10.1101/2024.10.25.620289

**Authors:** Eun Kim, Muhammad S. Khan, Juyeop Shin, Shaohua Huang, Alessandro Ferrari, Donghoon Han, Eunjin An, Thomas W. Kenniston, Irene Cassaniti, Fausto Baldanti, Dohyeon Jeong, Andrea Gambotto

## Abstract

COVID-19 vaccines effectively prevent symptomatic infection and severe disease, including hospitalization and death. However, unequal vaccine distribution during the pandemic, especially in low- and middle-income countries, has led to the emergence of vaccine-resistant strains. This underscores the need for alternative, safe, and thermostable vaccine platforms, such as dissolved microneedle array patches (MAP) delivering a subunit vaccine, which eliminate the need for cold chain and trained healthcare personnel. This study demonstrates that the SARS-CoV-2 S1 monomer with RS09, a TLR-4 agonist peptide, serves as an optimal protein subunit immunogen. This combination stimulates a stronger S1-specific immune response, resulting in binding to the membrane-bound spike on the cell surface and ACE2-binding inhibition, compared to the monomer S1 alone or trimer S1, regardless of RS09. MAP delivery of the rS1RS09 subunit vaccine elicited higher and longer-lasting immunity compared to conventional intramuscular injection. S1-specific IgG levels remained significantly elevated for up to 70 weeks post-administration. Additionally, different doses of 5, 15, and 45 *μ*g of MAP vaccines induced robust and sustained Th2-prevalent immune responses, suggesting a dose-sparing effect and inducing significantly high neutralizing antibodies against the Wuhan, Delta, and Omicron variants at 15 and 45 *μ*g dose. Moreover, gamma irradiation as a terminal sterilization method did not significantly affect immunogenicity, with irradiated vaccines maintaining comparable efficacy to non-irradiated ones. The stability of MAP vaccines was evaluated after long-term storage at room temperature and refrigeration for 19 months, showing minimal protein degradation. Further, after an additional one-month of storage at elevated temperature (42°C), rS1RS09 in both non-irradiated and irradiated MAP degraded less than 3%, while the liquid subunit vaccine degraded over 23%. Overall, these results indicate that gamma irradiation sterilized MAP-rS1RS09 vaccines maintain stability during extended storage without refrigeration, supporting their potential for mass production and widespread use in global vaccination efforts.

## 1. Introduction

The COVID-19 pandemic has left an indelible mark on the world, with over 775 million confirmed cases and nearly 7 million lives lost globally (until April 14, 2024) [1,2]. The United States alone has witnessed more than 1.18 million deaths [3]. Despite the pivotal role of vaccination efforts in combating the pandemic, challenges persist due to the ever-evolving nature of the SARS-CoV-2 virus, which continuously accumulates mutations in its genetic code. This dynamic leads to the emergence of new variants, some of which may evade immunity conferred by previous vaccinations, rising concern of the development of updated booster shots to address the evolving threat.

A critical issue during the pandemic is the unequal distribution of vaccines worldwide. Many low- to middle-income countries have struggled to secure adequate vaccine supplies [4,5], promoting the emergence of vaccine-resistant SARS-CoV-2 strains due to high infection rates in unvaccinated regions [6,7]. In response, alternative delivery methods such as dissolved microneedle array patches (MAP) for subunit vaccine administration are being developed and considered. Indeed, the MAP technology has recently been ranked as the highest global innovation priority for achieving equity of vaccine coverage in low- and middle-income countries by a consortium including the Gavi Secretariat, World Health Organization (WHO), Bill & Melinda Gates Foundation, UNICEF and PATH [8]. MAP delivery offers various advantages, including dose sparing, reduced needle-stick injuries, and the potential for painless self-administered injections, thus alleviating needle phobia in patients [9–12]. Moreover, MAP-based vaccines can be pre-formulated and stored stably for extended periods at room temperature, facilitating distribution even in regions with limited cold chain supply networks [13,14]. Additionally, MAP-based intradermal delivery has shown promising results in improving vaccine immunogenicity and safety in a number of vaccines studies of SARS-CoV-2 [15–20] and have been the subject of Phase I/II clinical trials of influenza, Japanese encephalitis, measles and rubella vaccination [12,21–24]. These clinical trials supported that the MAP vaccination is not only safe and well tolerated but also equal to or more effective than the conventional SC or IM injection.

Our prior findings have demonstrated the efficacy of MAP-based platforms in eliciting robust, long-lasting, and cross-neutralizing antibodies against SARS-CoV-2 variants, including Omicron and its subvariants [25]. This longevity in antibody response suggests that MAP-based vaccines could offer durable protection against SARS-CoV-2 and its variants. Therefore, the optimization of variant specific MAP boosters becomes essential to enhance vaccine effectiveness against current and future pandemics. To facilitate the clinical use of MAP platforms, terminal sterilization is a crucial consideration, albeit one that can impact the final product’s cost [26]. Various terminal sterilization methods have been developed, including dry-heat, steam, chemical sterilization (with ethylene, formaldehyde, or peracetic acid), and ionizing radiation (electron beam [e-beam], gamma ray [ɣ-ray], and X-ray) [27–29]. However, among them, gamma radiation of a minimum absorbed dose of 25 kGy is regarded as adequate for sterilizing pharmaceutical due to its ability to destroy microorganisms without heat, moisture, or chemical exposure [30]. However, concerns regarding the potential degradation of MAP morphology or the subunit vaccine within the polymer matrix due to high-energy transfer from gamma-ray irradiation must be addressed. First, we selected an ideal subunit vaccine by comparison the immunogenicity of the SARS-CoV-2 S1 protein depending on whether it was in monomeric or trimeric form, and in the presence of RS09, a TLR4 agonist. Following research aims to evaluate the immunogenicity of MAP delivery compared to conventional intramuscular injection, dose sparing effects, and long-term immunity of irradiated and non-irradiated MAP containing SARS-CoV-2 S1 subunit vaccine in BALB/c mice. Additionally, the stability of the SARS-CoV-2 S1 subunit vaccine in both irradiated and non-irradiated forms after long-term shelf storage at different temperatures (4°C, room temperature) was also assessed. Our findings suggest that immunogenicity remains comparable between irradiated and non-irradiated MAP, indicating the feasibility of long-term storage. These studies underscore the potential of gamma irradiation as a preferred method for the terminal sterilization of MAP at a commercial scale, enabling widespread distribution to combat the COVID-19 pandemic on a global scale.

## 2. Materials and methods

### 2.1. Construction of Recombinant Protein-expressing Plasmids

The coding sequence for the Delta (B.1.617.2) SARS-CoV-2-S1 glycoprotein mutated T19R; T95I; G142D; Del156-157; R158G; L452R; T478K; D614G, encompassing amino acids 1 to 661 of full-length from BetaCoV/Wuhan/IPBCAMS-WH-05/2020 (GISAID accession id. EPI_ISL_403928) [25,31] with C-tag (EPEA) flanked with Sal I and Not I sites was codon-optimized using UpGene algorithm for optimal expression in mammalian cells [32,33] and cloned into pAdlox. Similarly, for S1RS09, spanning amino acids 1 to 661 and equipped with the Bam HI−RS09 (APPHALS, TLR4 agonist)−EPEA, synthesis and cloning into pAdlox were performed using the same method [31]. For S1f and S1fRS09, RS09−EPEA in S1RS09 was replaced with bacteriophage T4 fibritin trimerization domain, foldon (f)−EPEA and fRS09−EPEA at Bam HI and Not I sites, respectively. pAd/S was generated by subcloning the codon-optimized SARS-CoV-2-S2 glycoprotein gene, amino acids 662 to 1273 of full-length spike, into the pAd/S1RS09 at BamH I/Not I sites. The plasmid constructs were confirmed by DNA sequencing.

### 2.2. Recombinant Proteins Expression and Purification

The production of SARS-CoV-2 Delta rS1, rS1RS09, rS1f, and rS1fRS09 involved transient expression in Expi293 cells with pAd/S1, pAd/S1RS09, pAd/S1f and pAd/S1fRS09, respectively, utilizing the ExpiFectamie^TM^ 293 Transfection Kit (ThermoFisher) as previously reported [25,31,34]. Subsequently, the recombinant proteins were purified using a CaptureSelect^TM^ C-tagXL Affinity Matrix prepacked column (ThermoFisher), followed the manufacturer’s guidelines as previously detailed [25,31,34]. In brief, the C-tagXL column underwent conditioning with 10 column volumes (CV) of equilibrate/wash buffer (20 mM Tris, pH 7.4) before sample application. The supernatant was adjusted to 20 mM Tris with 200 mM Tris (pH 7.4) before loading onto a 5-mL prepacked column at a rate of 5 ml/min. The column underwent subsequent washing cycles, alternating between 10 CV of equilibrate/wash buffer, 10 CV of strong wash buffer (20 mM Tris, 1 M NaCl, 0.05% Tween-20, pH 7.4), and 5 CV of equilibrate/wash buffer. The recombinant proteins were eluted from the column using an elution buffer (20 mM Tris, 2 M MgCl_2_, pH 7.4). The eluted solution was desalted and concentrated with phosphate buffered saline (PBS) in an Amicon Ultra centrifugal filter device with a 50,000 molecular weight cutoff (Millipore). The concentration of the purified recombinant proteins was determined by the BCA protein assay kit (Thermo Scientific) with bovine serum albumin (BSA) as a protein standard. The proteins were separated by reducing sodium dodecyl sulfate polyacrylamide gel electrophoresis (SDS-PAGE) and visualized through silver staining.

### 2.3. SDS-PAGE, Silver Staining, and Western Blot

The purified proteins were subjected to SDS-PAGE and visualized through silver staining and western blot. Briefly, after the supernatants were boiled in Laemmli sample buffer containing 2% SDS with beta-mercaptoethanol (β-ME), the proteins were separated by Tris-Glycine SDS-PAGE gels and transferred to nitrocellulose membrane. After blocking for 1 hour at room temperature (RT) with 5% non-fat milk in TBS-T, rabbit anti-SARS-CoV spike polyclonal antibody (1:3000) (Sino Biological) was added and incubated overnight at 4°C as primary antibody, and horseradish peroxidase (HRP)-conjugated goat anti-rabbit IgG (1:10000) (Jackson immunoresearch) was added and incubated at RT for 1 hours as secondary antibody. After washing, the signals were visualized using ECL Western blot substrate reagents and iBright 1500 (Thermo Fisher).

### 2.4. Preparation of Dissolving Microneedle Array Patches

Dissolving microneedle array patch containing the protein rS1RS09 were fabricated using the DEN (droplet extension technique) method. In this technique, droplets were solidified and formed into cone shape microstructure by air blowing, as previously descried [25,35]. The MAP used in this study was provided by Raphas Co., Ltd. (Rep. of Korea) and fabricated to three different forms changing the array number and patch size depending on dosage. Briefly, for MAP-rS1(WU+Beta), 35 base arrays in 1.86 cm X 1.48 cm (2.75 cm^2^) for 7 μg of patch and 25 base array in 1.6 cm diameter (2.0 cm^2^) for 5 μg of patch, respectively, were fabricated. For dose sparing of MAP-rS1RS09, 35 base arrays in 1.86 cm X 1.48 cm (2.75 cm^2^) for 45 μg of patch, 12 and 4 base arrays in 1.6 cm diameter (2.0 cm^2^) for 15 and 5 μg of patches, respectively, were fabricated. All patches were dispensed onto a pattern-mask hydrocolloid adhesive sheet (Hiks C&T) using a customized MPP-1 dispenser (Musashi) with 10% hyaluronic acid (HA, Kikkoman). The dispensed array was then dried overnight at RT. Subsequently, a viscous solution containing the antigens and HA (Contipro Inc) was dispensed onto a different base array. The two dispensed droplets were brought into contact and extended to the target length, and then symmetric air blow was applied at RT to solidify the extended viscous droplets, forming cone-shaped microstructures. The mechanical strength of each MAP was measured using the universal test machine (UTM, ZwickRoell). Briefly, a MAP was mounted on the UTM stage and aligned with the UTM probe. The probe was moved vertically until the microneedle fractured, with the force responsible for breaking the microneedle recorded as its mechanical strength. For final packaging, each MAP was placed in a polyethylene terephthalate (PET) blister pack ; the blister pack was sealed in light-protective foil pouches (thickness: 100 μm) with desiccant. In line with our efforts toward clinical production of MAP-rS1RS09Delta, we sterilized a small group of these MAPs using gamma irradiation to determine clinically relevant sterilizing conditions and any effect on immunogenicity. Gamma-ray irradiation was conducted using a cobalt-60 irradiator (JS-8900, Nordion Inc.) at 15∼30 kGy.

### 2.5. Animal Immunization

Female BALB/c mice (n=5 animals per group) were bled from the retro-orbital vein before immunization and primed with 45*μ*g of either Delta rS1, rS1RS09, rS1f, or rS1fRS09. Mice were bled on week 3 and received a homologous booster of 45*μ*g of each protein. Subsequent bleeds were performed on weeks 5, 7, and 18. For the study of long-term immunogenicity, BALB/c mice (5 per group) were primed and boosted at three-week intervals with 45*μ*g of rS1RS09 intramuscularly. Serum samples were collected in weeks 0, 3, 5, 7, 9, 12, 16, 20, 28, 40, 71, 90, and 104 after prime immunization.

In the experiment with ICR mice, 5 *μ*g of rS1(Wu+Beta) were injected into the thigh intramuscularly, and MAPs loaded with either 5 or 7 *μ*g of rS1(Wu+Beta) were applied to the skin of the back region of female ICR mice (n = 5 animals per group). Prior to vaccination, the hair at the vaccination site was removed by shaving and depilatory cream. MAPs were applied to the dehaired back skin of the mice using a handheld spring applicator and held in place by finger pressure for 10 seconds, followed by leaving it on the skin for more than 2 hours. No adverse skin effects, such as skin irritation at the vaccinated region, were observed. At 3 weeks after the primary immunization, mice received booster immunizations with homologous immunogens. Blood samples were collected from the retro-orbital vein of mice every three weeks until week 9. The obtained serum samples were diluted and used to evaluate rS1-specific antibodies by ELISA.

MAPs loaded with 5, 15, and 45 *μ*g of Delta rS1RS09, referred to as MAP-rS1RS09 with or without irradiation, were applied to the skin of the back region of female BALB/c mice (n = 5 animals per group). MAP application was performed using the same method as with ICR mice. At 3 weeks after the primary immunization, mice were received booster immunizations with homologous immunogens. Blood samples were collected from the retro-orbital vein of mice every three to ten weeks until week 104. The obtained serum samples were diluted and used to evaluate rS1-specific antibodies by ELISA and VNT assay. Since aged mice develop spontaneous leukemias and other tumors, the dedicated veterinarians oversee the animals’ physical and psychological health and ruled out any mice having diseases that may influence immune responses. Indeed, one mouse of 15 *μ*g non-irradiated MAP-rS1RS09 group was ruled out at week 71, and two to three mice were ruled out in week 90, and two to four mice in week 104, because they were euthanized due to tumors or were found deceased. Mice were maintained under specific pathogen-free conditions at the University of Pittsburgh, and all experiments were conducted in accordance with animal use guidelines and protocols approved by the University of Pittsburgh’s Institutional Animal Care and Use (IACUC) Committee.

### 2.6. Assessment of Serum Humoral Antibodies

Serum humoral antibodies generated against spike protein were assessed using ELISA, as previously described [32,36]. Sera from all mice were collected before vaccination and then every 2 to 10 weeks after immunization. These sera were tested for SARS-CoV-2 S1WU-specific IgG antibodies using conventional ELISA. Furthermore, sera collected at weeks 0, 9, 28, 52, and 71 or at weeks 0, 5/6, 20, 40, and 71 after vaccination were also examined for SARS-CoV-2-S1-specific IgG1 and IgG2a antibodies using ELISA. In brief, ELISA plates were coated with 200 ng of recombinant SARS-CoV-2-S1WU protein (Sino Biological) or house purified rS1WU per well overnight at 4°C in carbonate coating buffer (pH 9.5) and then blocked with PBS containing 0.05% Tween 20 (PBS-T) and 2% BSA for one hour. Mouse sera were serially diluted in PBS-T with 1% BSA and incubated overnight. After washing the plates, anti-mouse IgG-horseradish peroxidase (HRP) (1:10000, Jackson Immunoresearch) was added to each well and incubated for one hour. For detection of IgG1 and IgG2a, HRP-conjugated anti-mouse IgG1 and IgG2a (1:20000, Jackson Immunoresearch) were added to each well and incubated for 1 hour. The plates were washed three times, developed with 3,3’5,5’-tetramethylbenzidine, and the reaction was stopped. Optical densities (ODs) were read at 450 nm with a SpectraMax iD5 microplate reader (Molecular Devices).

### 2.7. Flow Cytometry

Two weeks after the booster immunization, pooled sera were obtained from all mice and screened for SARS-CoV-2-S-specific antibodies using fluorescence-activated cell sorter (FACS) analysis of Expi293 cells transfected with either pAd/S or pAd control using ExpiFectamie^TM^ 293 as described previously. Briefly, at 36 hours after transfection, the cells were harvested, counted, and washed with PBS. Half a million cells were incubated with 1 *μ*l of mouse serum from each group or 1 *μ*l of monoclonal antibody (D003, Sino Biological) for 30 min, followed by staining with a FITC-conjugated anti-mouse secondary IgG antibody (Jackson ImmunoResearch). Data acquisition and analysis were performed using LSRII (BD). The results were calculated as follows: Positive cells (%) = 100 × (positive cell (%) using each mouse serum / positive cell (%) using monoclonal antibody).

### 2.8. ACE2 Blocking Assay

Antibodies blocking the binding of SARS-CoV-2 spike including Wuhan, Omicron (BA.1), Omicron sub-variants (BA.2, BA.3, BA.1+R346K, BA.1+L452R), Delta lineage (AY.4), Alpha (B.1.1.7), Beta (B.1.351), and France (B.1.640.2) to ACE2 were detected with a V-PLEX SARS-CoV-2 Panel 25 (ACE2) Kit (Meso Scale Discovery (MSD)) according to the manufacturer’s instructions. The assay plate was blocked for 30 min and washed. Serum samples were diluted (1:25 or 1:100) and 25 μl were transferred to each well. The plate was then incubated at RT for 60 min with shaking at 700 rpm, followed by the addition of SULFO-TAG conjugated ACE2, and continued incubation with shaking for 60 min. The plate was washed, 150 μl MSD GOLD Read Buffer B was added to each well, and the plate was read using the QuickPlex SQ 120 Imager. Electrochemiluminescent values (ECL) were generated for each sample. The lower values than pre-immunized sera were adjusted with the values at week 0. Results were calculated as % inhibition compared to the negative control for the ACE2 inhibition assay, and % inhibition is calculated as follows: % neutralization = 100 × (1 − (sample signal/negative control signal)).

### 2.9. SARS-CoV-2 microneutralization assay

Neutralizing antibody titers against SARS-CoV-2 were defined according to the following protocol (59, 60). Briefly, 50 µl of sample from each mouse, starting from 1:10 in a twofold dilution, were added in two wells of a flat bottom tissue culture microtiter plate (COSTAR), mixed with an equal volume of 100 TCID_50_ of a SARS-CoV-2 Wuhan, Delta (B.1.617.2), or Omicron (BA.1) strain isolated from symptomatic patients, previously titrated. After 1 hour incubation at 33°C in 5% CO_2_, 3 x 10^4^ VERO E6 cells were added to each well. After 72 h of incubation at 33°C 5% CO_2_, wells were stained with Gram’s crystal violet solution plus 5% formaldehyde 40% m/v (Carlo ErbaSpA, Arese, Italy) for 30 min. After washing, wells were scored to evaluate the degree of cytopathic effect (CPE) compared to the virus control. Neutralizing titer was the maximum dilution with a reduction of 90% of CPE. A positive titer was equal to or greater than 1:10. The GMT of VNT_90_ endpoint titer was calculated with 5 as a negative shown <1:10. Sera from mice before vaccine administration were always included in VNT assay as a negative control.

### 2.10. MAP Storage and Reconstitution for Analysis

The packed MAPs, stored in light-protective pouches with desiccant, were kept in a refrigerator at 4 °C or in a drawer at room temperature (RT) for one week or nineteen months. To assess long-term stability, the packed MAPs stored at 4 °C for nineteen months were further stored for one month in a refrigerator at 4 °C, in a drawer at RT, and in temperature-controlled incubator at 42 °C to represent storage conditions in a hot climate. MAPs were removed from the pouches and placed in each well of a 6-well plate with 2 ml of PBS. Reconstitution was achieved by incubating for 30 min at RT with shaking at 700 rpm. The resulting solution containing rS1RS09 and microneedle matrix excipients was transferred to a centrifuge tube and stored at −80 °C until further analysis. The stability assay was performed using SDS-PAGE and silver staining, followed by analysis using ImageJ v1.47 software (National Institutes of Health, Bethesda, MD, USA) to determine the relative intensity of each protein band for comparison. The results are presented as the percentage of degradation proteins relative to the smear ratio, which is calculated by dividing the integrated density of the sample band area by that of all protein bands in the preparation lane. The percentage of recombinant protein degradation in the reconstructed MAP or in the buffer was calculated as follows: % protein degradation = 100 × [(smear ratio of negative control – smear ratio of sample MAP)/smear ratio of negative control]. The negative control comprised non-irradiated MAP stored at 4°C or recombinant proteins stored at -20°C for 19 months. To compare the stability of rS1RS09 in the buffer solution, recombinant proteins (rP) stored for 19 months at –20°C were 10-fold diluted in PBS or in the preservative buffer (10mM Tris, 75mM NaCl, 2mM MgCl_2_, 5% Trehalose), divided into two microtubes, and further stored for one month at –20°C or 42°C in a humidity chamber. The percentage of recombinant protein degradation was calculated in the same way as above.

### 2.11. Statistical analysis

Statistical analyses were performed using GraphPad Prism v10 (San Diego, CA). Antibody endpoint titers and neutralization data were analyzed by Kruskal-Wallis test, followed by Dunn’s multiple comparisons. A Mann-Whitney U test was used for intergroup statistical comparison. Comparison of intramuscular injection and MAP groups were analyzed by Bartlett’s test. Data were presented as the means ± standard errors of the mean (SEM) or geometric means ± geometric errors. Significant differences are indicated by ∗ p < 0.05, ∗∗p < 0.01, ∗∗∗p < 0.001.

## 3. Results

### 3.1. Purification of Recombinant Proteins

Initially, we compared the immunogenicity of the SARS-CoV-2 S1 protein in monomeric versus trimeric forms, and in the presence of RS09, a TLR4 agonist. To produce four recombinant proteins of the Delta SARS-CoV-2-S1, both in the absence and presence of the bacteriophage T4 trimer domain foldon (f) and RS09, we generated pAd/S1, pAd/S1RS09, pAd/S1f, and pAd/S1fRS09 by subcloning the codon-optimized Delta SARS-CoV-2-S1 gene with a C-tag (EPEA) into the shuttle vector pAd (GenBank U62024) at the *Sal* I and *Not* I sites (**Fig. 1A**). Variant-specific mutations for SARS-CoV-2 Delta (B.1.617.2) S1 proteins are outlined. For the expression of the four Delta S1 subunit proteins, Expi293 cells were transfected with the plasmids pAd/S1, pAd/S1RS09, pAd/S1f, and pAd/S1fRS09. Five days after transfection, the supernatants of Expi293 cells were collected, and the four recombinant proteins (Delta rS1, rS1RS09, rS1f, and rS1fRS09) were purified using C-tagXL affinity matrix and confirmed by silver staining and Western blot (**Fig. 1B and 1C**). Each S1 recombinant protein was recognized by a polyclonal anti-spike of SARS-CoV-2 antibody at the expected glycosylated monomeric molecular weights of about 110 kDa under denaturing reduced conditions. These results indicates that four recombinant proteins (Delta rS1, rS1RS09, rS1f, and rS1fRS09) were successfully purified for subsequent experiments.

**Fig. 1.**
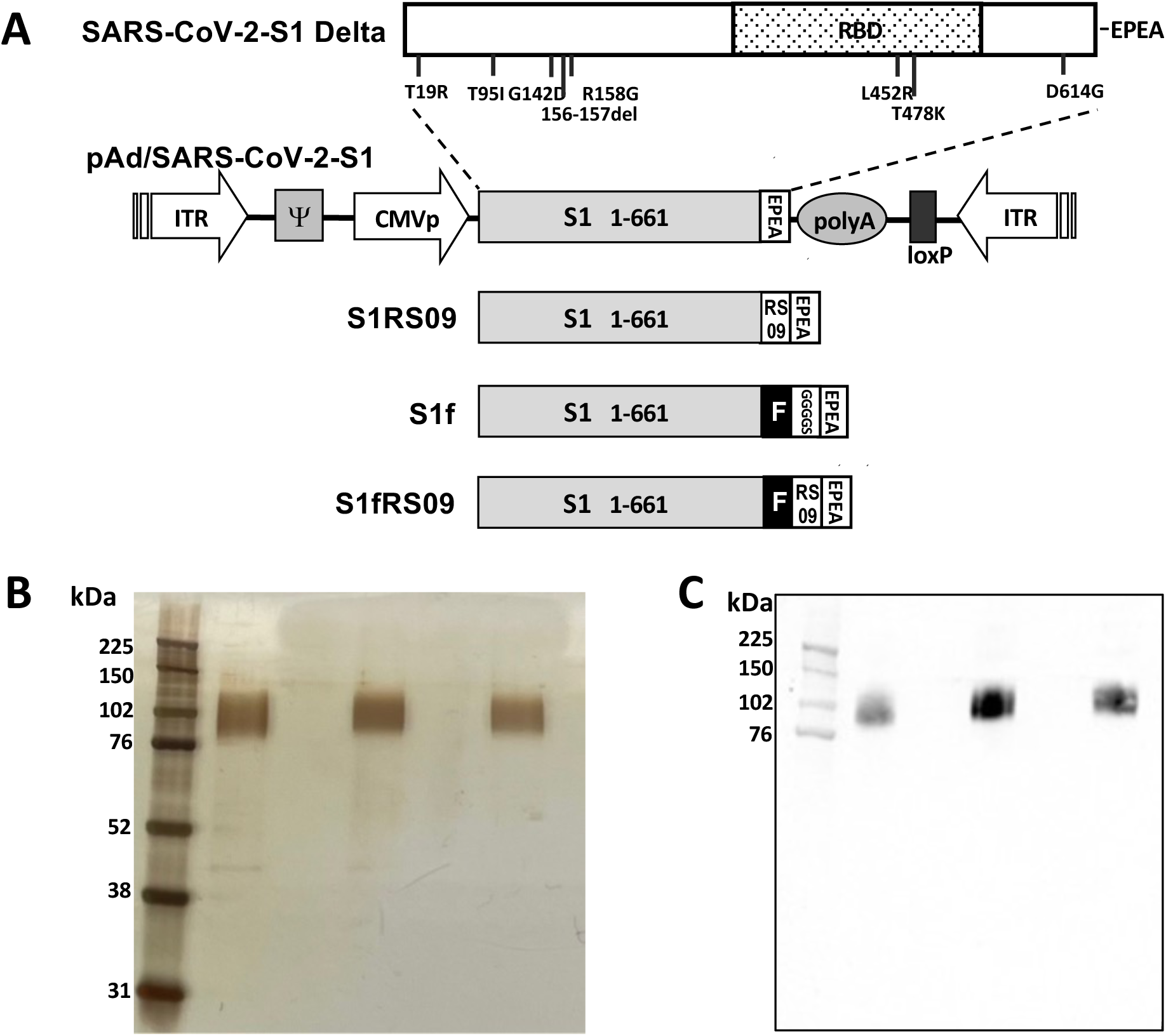
Construction and expression of recombinant SARS-CoV-2-S1Delta proteins. (A) A shuttle vector carrying the codon-optimized four constructs of SARS-CoV-2-S1Delta gene encoding N-terminal 1-661 with c-tag (EPEA, glutamic acid-proline-glutamic acid-alanine) was designated as shown in the diagram. Amino acid changes in the SARS-CoV-2-S1 region of in this study are shown. ITR: inverted terminal repeat; CMVp, cytomegalovirus promoter; RBD: receptor binding domain. (B) Purified proteins, rS1RS09 (lane1), rS1f (lane2), and rS1fRS09 (lane3), isolated by c-tag affinity purification were separated by SDS-PAGE and visualized by silver staining. Molecular weight marker (MW marker) is indicated on the left. (C) Detection of the Purified SARS-CoV-2-S1 proteins, rS1RS09 (lane1), rS1f (lane2), and rS1fRS09 (lane3), respectively, by western blot using rabbit anti spike of SARS-CoV Wuhan polyclonal antibody.

### 3.2. Identification of S1RS09 as an optimal immunogen

To compare the immunogenicity of four Delta S1 proteins, BALB/c mice (6–8 weeks old, n = 5) were immunized intramuscularly (IM) with 45 *μ*g of either Delta rS1, rS1RS09, rS1f, or rS1fRS09 in a prime-boost regimen, spaced three weeks apart. Blood samples were collected before the initial immunization and at weeks 3, 5, 7, and 18, and then examined for the presence of antibodies specific to SARS-CoV-2-S1 using ELISA **(Fig. 2A and 2B)**. As shown in Fig. 2B, statistically significant S1-specific IgG EPT was induced in the mice immunized with either S1 or S1RS09 at weeks 5, 7 and 18, while the mice vaccinated with trimers, rS1f or rS1fRS09 showed lower S1-specific IgG EPT. Interestingly, more statistically significant S1-specific IgG EPT was induced in the mice immunized with S1RS09 all time points (**Fig. 2B**). These findings suggest that SARS-CoV-2 S1 monomer in the presence of RS09 (rS1RS09) is more immunogenic than the trimer S1 subunit protein regardless of RS09.

**Fig. 2.**
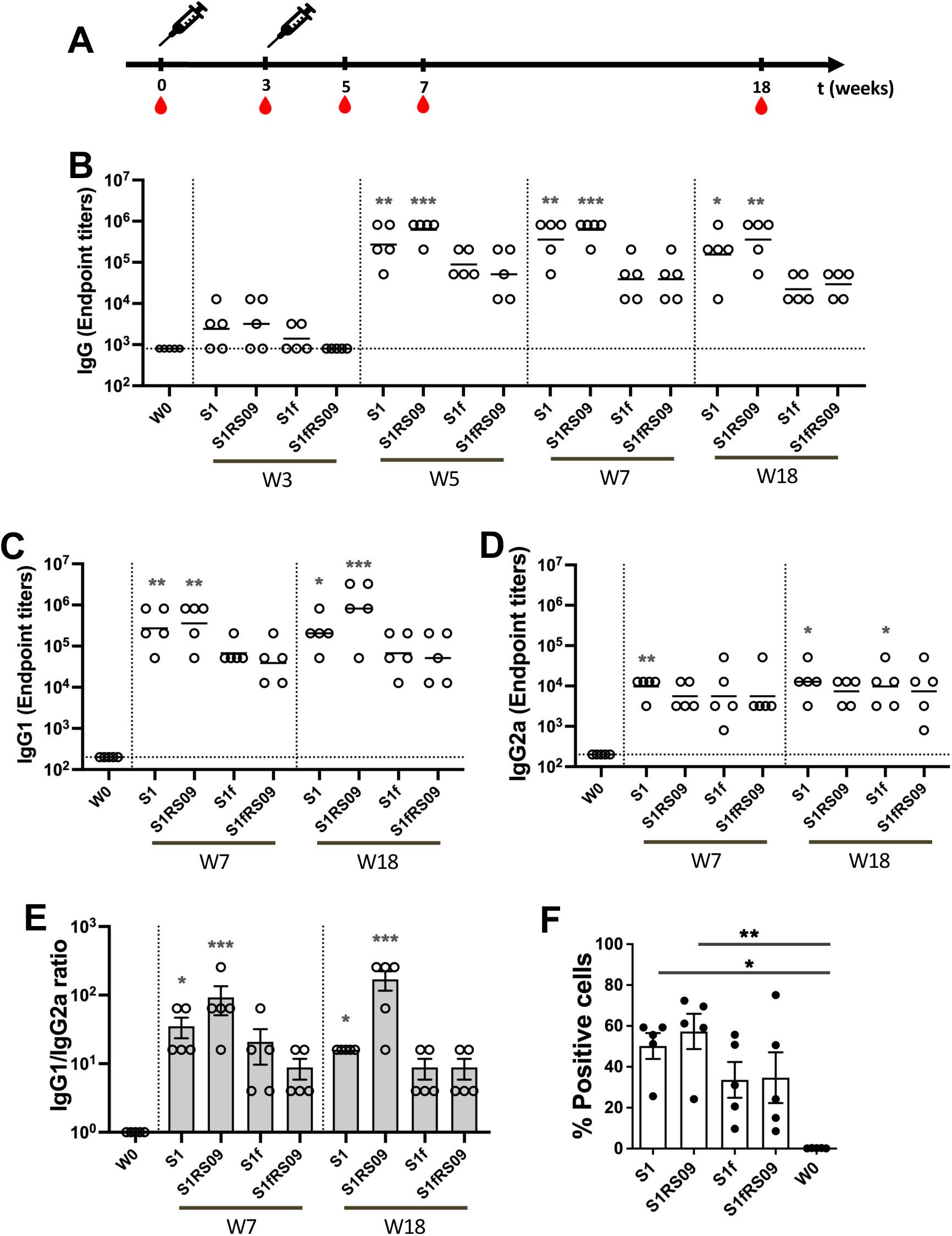
Comparison in mouse immunized with four SARS-CoV-2 rS1Delta protein subunit vaccines. (A) Schedule of immunization and blood sampling for IgG end point titration. Balb/c mice (5 per group) were immunized with 45μg of rS1, rS1RS09, rS1f, and rS1fRS09 proteins of SARS-CoV-2 Delta then administered intramuscularly at week 0 and 3. Syringes indicated the timing of immunizations, and the red drops denote times at which blood was drawn. (B) Sera were diluted, and SARS-CoV-2-S1-specific antibodies were quantified by ELISA to determine the IgG endpoint titer. The IgG titers at each time points were showed in each mouse. The bars represent geometric mean with geometric SD. (C and D) Sera at weeks 0, 5, and 18 were diluted, and SARS-CoV-2-S1WU-specific IgG1 (**C**) an IgG2a (**D**) were quantified by ELISA to determine each IgG subclasses endpoint titer. The titers at each time points were showed for each mouse. The bars represent geometric mean. (E) S1-specific IgG1/IgG2a ratios of individual mice at weeks 7 and 18 as mean values with SEM. (F) Flow cytometry assay of Expi293 cells expressing S1Delta-S2WU at the cell surface. At 36 hrs post-transfection with pAd/ S1Delta-S2WU, binding to SARS-CoV-2-S at the Expi293 cell surface was analyzed by incubation with mice sera obtained at week 7 after immunization followed by staining with FITC-conjugated anti-mouse IgG. Groups were compared by the Kruskal-Wallis test at each time point, followed by Dunn’s multiple comparisons. Significant differences are indicated by *p < 0.05, **p < 0.01, ***p < 0.001, n.s., not significant.

We also examined the S1-specific IgG isotypes switch, IgG1 and IgG2a following immunization with the four Delta S1 protein subunits, indicating a prevalent and/or balanced Th2- or Th1-biased immune response, respectively (**Fig. 2C and D**). Sera collected at weeks 0, 7, and 18 post-prime were subjected to isotype-specific ELISA. The S1-specific IgG1 showed a significant increase in the mice immunized with either S1 or S1RS09 at week 7 and maintained significantly high levels until week 18 comparison to pre-immunized sera, although the geometric mean titer (GMT) diverged slightly lower in the S1 group and slightly higher in the S1RS09 group (**Fig. 2C)**. In case of S1-specific IgG2a, only S1-immunized mice showed a significant increase at week 7, whereas significant increases were detected in the mice immunized either S1 or S1f at week 18. Interestingly, the GMT of IgG2a increased slightly at week 18 in all groups compared to those at week 7 (**Fig. 2D**). Th2-prevalent responses were observed at all time points based on the ratio of IgG1 to IgG2a antibody subclasses in all the groups at week 7, with slightly higher ratios in the S1RS09 group. It was observed that the S1RS09 group induced more toward Th2-biased immune response by higher IgG1 and similar IgG2a at week 18, while the three other groups switched slightly toward a Th1 response or remained steady (**Fig. 2E**). The ratio of IgG1/IgG2a at week 18 was approximately 10-fold higher when the vaccine was administered with S1RS09 compared to the S1 group.

Next, we examined whether these antibodies could bind to membrane-bound full-length spike protein by measuring the reactivity on Expi293 cells transfected with pAd/S (S1Delta + S2Wuhan) or pAd using flow cytometry. The mice immunized with either Delta rS1, rS1RS09, rS1f, or rS1fRS09 developed membrane-bound S-specific antibodies, while no specific antibody response was detected in pre-immunized sera (**Fig. 2F**). The mean of percentage of positive cells was slightly higher in the rS1RS09-immunzed mouse group compared to other groups. No antibodies were detected on cells transfected with pAd (data not shown).

To assess the capacity of these vaccines to potentially neutralize RBD-ACE2 binding, we evaluated the ability of antibodies in the serum at week 7 to inhibit the binding between ACE2 and the trimeric spike protein of SARS-CoV-2 variants, representing a sensitive measure of neutralizing activity. We used V-PLEX SARS-CoV-2 (ACE2) Kit Panel 18, which included Wuhan, Alpha (B.1.1.7), Beta (B.1.351), Gamma (P.1), Delta (B.1.617.2), Zeta (P.2), Kappa (B.1.617.1), New York (B.1.516.1), India (B.1.617 and B.1.617.3) variants. The ability of antibodies to neutralize the interaction between spike protein of SARS-CoV-2 variants and ACE2 was examined in all animals immunized either rS1, rS1RS09, rS1f, or rS1fRS09 at weeks 0 and 7 post-prime at a dilution of 1:20. The ACE2 inhibitory activities of the sera from the mice immunized with Delta rS1, rS1RS09, rS1f, or rS1fRS09 against all variants were on average 50.9% ± 5.64, 57.1% ± 12.21, 32.4% ± 7.42, and 35.4% ± 5.53 at weeks 7, respectively, with 21.2% ± 7.15 at week 0. Overall, the median percent inhibition of the sera from rS1RS09-immunized groups was the highest against all variants among the groups, except against Beta and Gamma, which showed the highest inhibition in rS1-immunized group. Interestingly, the inhibitions against Wuhan, Alpha, Kappa, and India (B.1.617.3) spike proteins were significantly different compared to week 0 in rS1-immunized mice. The ACE2-binding inhibitions of rS1RS09-immunized mice were significantly increased for all tested SARS-CoV-2 variants, except Beta and Gamma when compared to week 0 (**Fig. 3**). They demonstrated moderate ACE2-binding inhibition from rS1f- and rS1fRS09-immunized sera, which were not statistically significant against all variants when compared to week 0. These findings suggest that SARS-CoV-2 S1 monomer in the presence of RS09 (rS1RS09) is more immunogenic than the trimer S1 subunit protein, regardless of RS09, considering it as an ideal vaccine candidate.

**Fig. 3.**
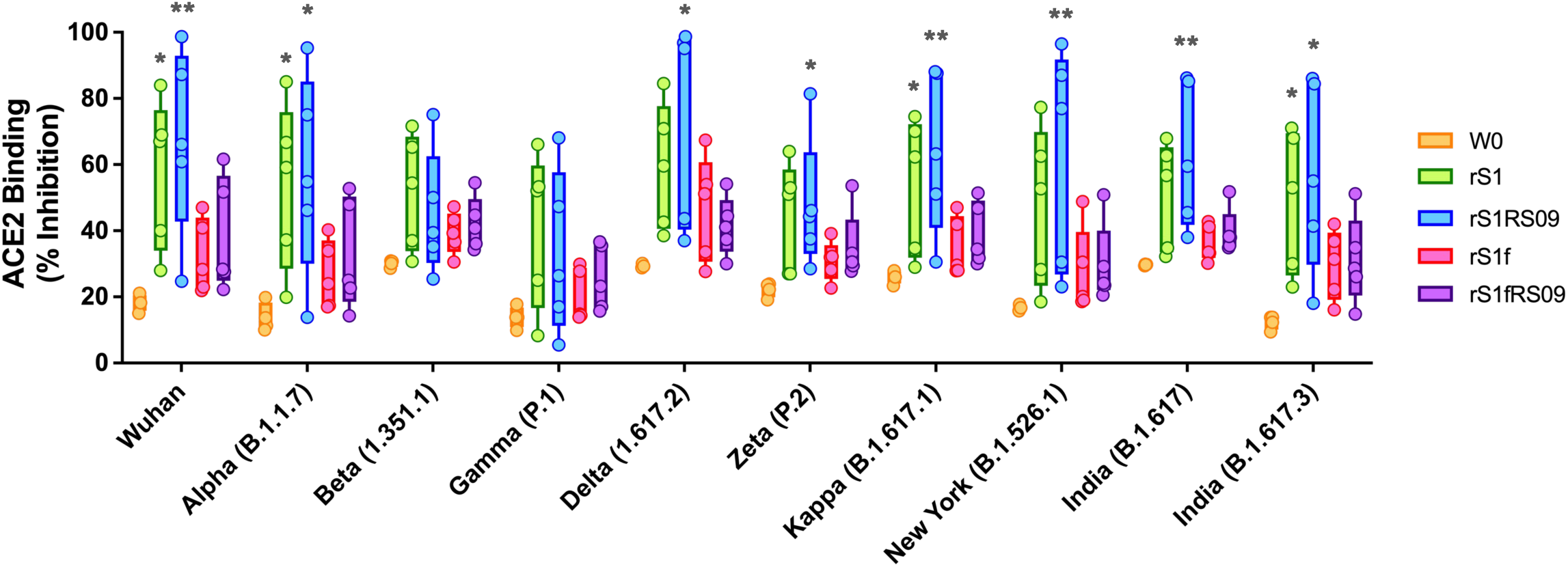
Percent ACE binding inhibition of neutralizing antibodies against SARS-CoV-2 variants. Antibodies in sera capable of neutralizing the interaction between SARS-CoV-2 Wuhan, Alpha (B.1.1.7), Beta (B.1.351), Gamma (P.1), Delta (B.1.617.2), Zeta (P.2), Kappa (B.1.617.1), New York (B.1.516.1), India (B.1.617 and B.1.617.3) variants spike and ACE2 were examined at week 0 (peach), and at week 7 post-prime in all animals immunized with rS1 (green), rS1RS09 (blue), rS1f (pink), and rS1fRS09 (purple) of SARS-CoV-2 Delta. Serum samples were diluted in 1:20 before adding the V-PLEX plates. Box and whisker plots represent the median and upper and lower quartile (box) with min and max (whiskers). Groups were compared by Kruskal-Wallis test at each variant, followed by Dunn’s multiple comparisons. Significant differences are indicated by *; p < 0.05 and **; p < 0.01. Asterisks represent statistical differences compared with pre-immunized sera (W0).

### 3.3. Enhanced Immunity of MAP delivery compared to conventional intramuscular injection

In a previous investigation, we demonstrated the effectiveness of a prime-boost SARS-CoV-2 S1 subunit MAP vaccine in eliciting specific antibody responses in mice [25]. We also compared the antibody responses induced by MAP delivered S1 subunit vaccine to those elicited by conventional intramuscular (IM) injections. Eight-week-old ICR mice were immunized with 5 *μ*g of rS1(Wu+Beta) (Group 1) intramuscularly (IM) into the thigh, while MAPs loaded with either 5 or 7 *μ*g of rS1(Wu+Beta) (Group 2 and 3, respectively) were inoculated intradermally (ID) three weeks apart (**Fig. 4A and 4B**). We measured S1-specific total IgG antibody endpoint titer (EPT) at weeks 3, 6, and 9 after the initial vaccination by ELISA. S1-specific antibodies in mice immunized with MAP-S1(Wu+Beta) (Group 2 and 3) showed significantly elevated IgG levels at weeks 6 and 9 following the booster vaccination compared to pre-immunized controls, whereas antibody levels in sera from IM-immunized mice (Group 1) were not significantly elevated compared to pre-immunized sera and declined slightly at week 9 (**Fig. 4C**). Furthermore, we examined the S1-specific IgG1 and IgG2a titers at week 6 to determine if there were significant differences due to the vaccine dose and route of immunization. IgG1 endpoint titers of Group 2 and Group 3 were statistically significant when compared to pre-immunized controls, while only Group 3 IgG2a endpoint titers were statistically significant (**Fig. 4D and 4E**). However, similar Th2-prevalent responses were observed regardless vaccine dose and route of immunization based on the ratio of IgG1 to IgG2a antibodies in all the groups (**Fig. 4F**). These findings suggest that the delivery of protein subunit vaccine via MAP is superior to conventional intramuscular injection, although similar Th2-prevalent responses are observed.

**Fig. 4.**
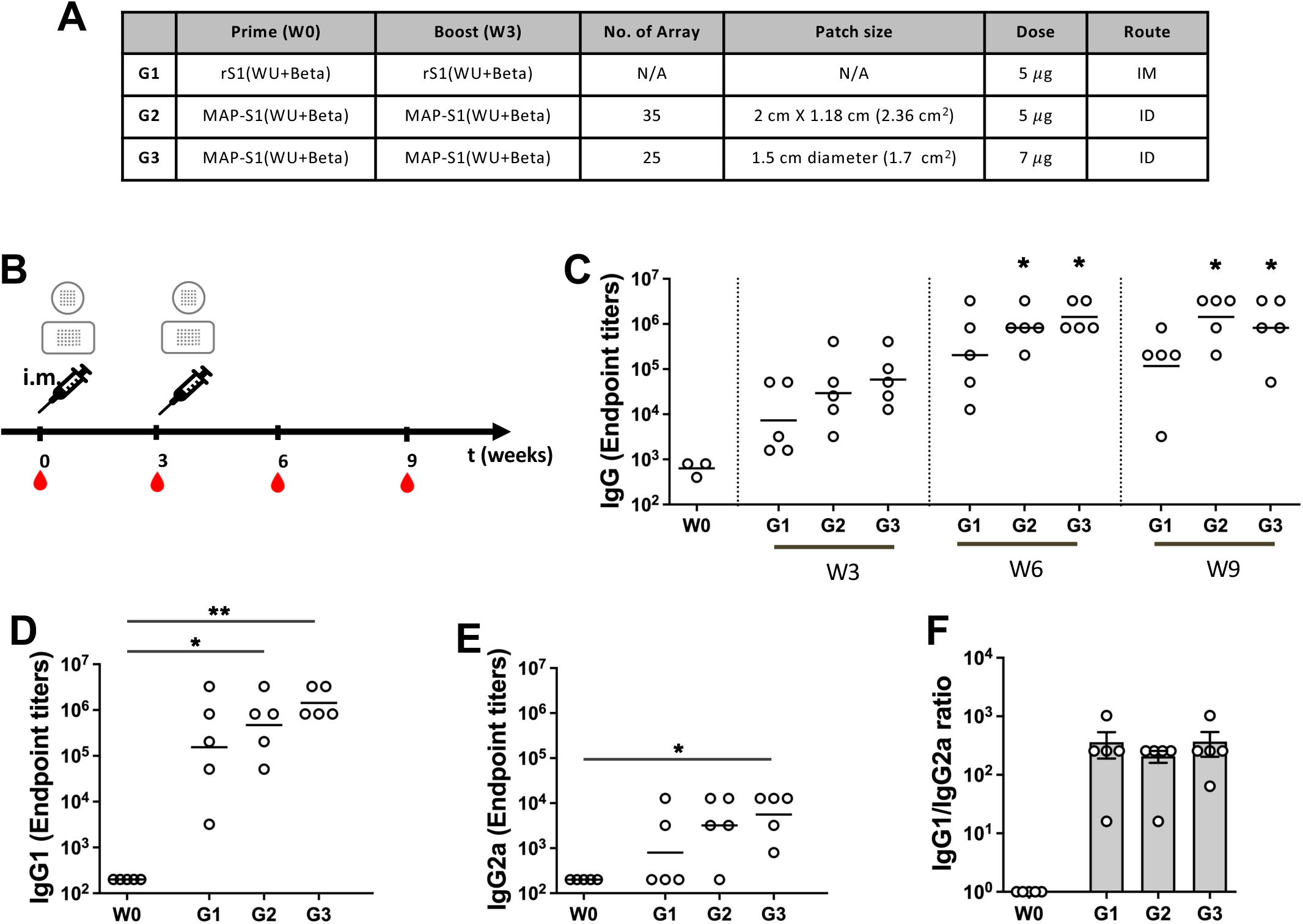
Immune responses induced by different vaccine routes of administration in ICR mice. (A) Immunogens and delivery routes of each group (B) Experimental schedule representing the immunization timeline. ICR mice (5 per group) were prime and boosted three weeks intervals with either 5*μ*g of rS1RS09Delta intramuscularly (IM) or MAP-rS1RS09Delta intradermally (ID). The red drops represented bleeding. (C) Reciprocal serum endpoint dilutions of SARS-CoV-2-S1-specific antibodies were measured by ELISA to determine the IgG endpoint titers at weeks 0, 3, 6, and 9. Sera at weeks 0 and 6 were diluted, and SARS-CoV-2-S1WU-specific IgG1 (D) an IgG2a (E) were quantified by ELISA to determine each IgG subclasses endpoint titer. The bars represent geometric mean. (F) S1-specific IgG1/IgG2a ratios of individual mice at weeks 6 as mean values with SEM. IM and MAP groups were compared for statistically significant differences using Kruskal-Wallis test at each variant, followed by Dunn’s multiple comparisons. *: p value,0.05; **: p,0.01.

### 3.4. Long-lasting Immunity of MAP delivery compared to conventional intramuscular injection

In a previous study, we demonstrated that a prime-boost SARS-CoV-2 S1 subunit MAP vaccine was more effective in triggering specific antibody responses in mice compared to intramuscular injection. To further evaluate the differences due to the vaccine dose and longevity, we generated and purified large-scale of the S1RS09 recombinant proteins and fabricated MAPs loaded 45 μg of rS1RS09 (MAP-rS1RS09) using a hyaluronic acid-based droplet extension technique (DEN), which offers the benefits of being faster, scalable, and cost-effective (**Fig. 5A**). The DEN MAP features a rectangle design with dimensions of 2 cm x 1.18 cm, housing a grid of 7×5 microneedle arrays, each measuring approximately 700 μm in length. The MAP consists of a double layered form with a base layer measuring 900 μm in base width and 200 ± 20 μm in height, and an upper layer measuring 500 ± 20 μm in length. Each MAP occupied an area size of 2.36 cm^2^, with 7 × 5 arrays of approximately 700 ± 50 μm total length (**Fig. 5B and 5C**). The mechanical strength (fracture force) of the MAP was measured to be more than 0.87 N, which is the required strength to penetrate the skin.

**Fig. 5.**
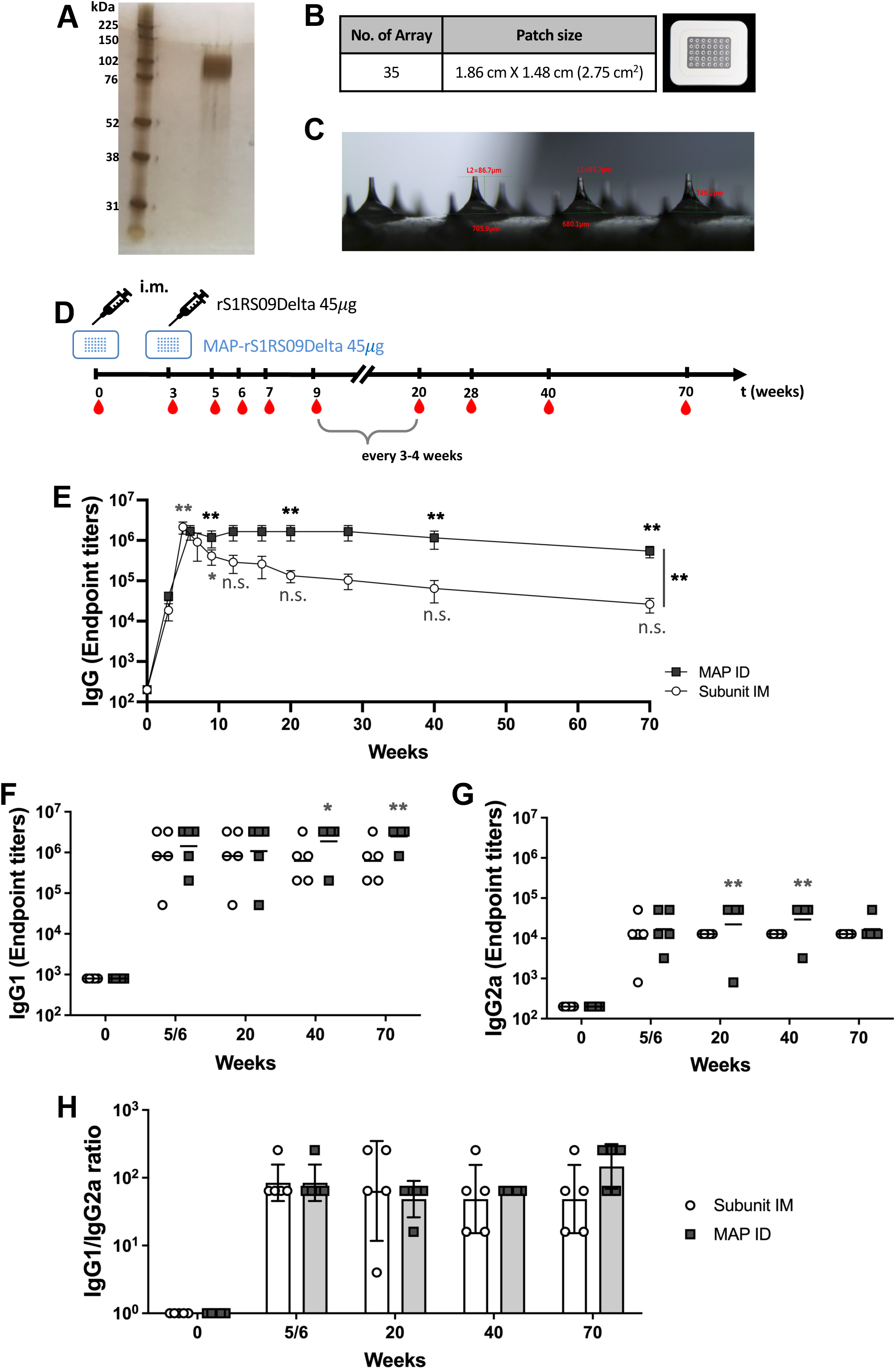
Immune responses induced by different vaccine routes of administration of high dose. (A) Silver stained SDS-PAGE of purified SARS-CoV-2 Delta rS1RS09 protein (B) Fabrication of dissolving microneedle patch, MAP-rS1RS09 was composed of 35 microneedles. (C) Each MN was approximately 700 *μ*m in length, occupying a 2.36 cm^2^ area size of patch, 7 × 5 arrays of 700 μm-long microneedles with 45*μ*g of rS1RS09Delta (D) Experimental schedule representing the immunization timeline. Balb/c mice (5 per group) were prime and boosted three weeks intervals with either 45*μ*g of rS1RS09Delta intramuscularly or MAP-rS1RS09Delta intracutaneously. The red drops represented bleeding. (E) Reciprocal serum endpoint dilutions of SARS-CoV-2-S1-specific antibodies were measured by ELISA to determine the IgG endpoint titers until week 90. Sera at weeks 0, 5, and 18 were diluted, and SARS-CoV-2-S1WU-specific IgG1 (F) an IgG2a (G) were quantified by ELISA to determine each IgG subclasses endpoint titer. The titers at each time points were showed for each mouse. The bars represent geometric mean. (H) S1-specific IgG1/IgG2a ratios of individual mice at weeks 7 and 18 as mean values with SEM. IM and MAP groups were compared for statistically significant differences using non-parametric Mann-Whitney-U-test compared with pre-immunized sera. *p < 0.05, **p < 0.01, ***p < 0.001, n.s., not significant.

To compare the long-term immunogenicity in the two different routes, BALB/c mice (5 per group) were primed and boosted at three-week intervals with either 45μg of rS1RS09 intramuscularly or MAP-rS1RS09 intradermally. Serum samples were collected at weeks 0, 3, 5/6, 7, 9, 12, 16, 20, 28, 40, and 70 after prime immunization (**Fig. 5D**). Serum samples were serially diluted to determine SARS-CoV-2-S1-specific IgG titers using ELISA. All vaccinated groups had significantly high and similar geometric mean S1 IgG antibody endpoint titer (EPT) at weeks 5 and 6 when compared to week 0, illustrating the superior immunogenicity conferred by boost immunization. The IgG geometric mean in mice immunized with MAP-rS1RS09 reached a steady geometric mean S1 IgG EPT through week 40, as high as the peak at week 6, and waned slightly thereafter. In contrast, mice immunized with rS1RS09 IM peaked at week 5, similar to the MAP group, but the immune response steadily waned through week 70, with significant difference between the two groups (p<0.01, Barttlet’s test) (**Fig. 5E**). S1-specific antibodies in mice immunized with MAP-rS1RS09 remained significantly elevated compared to pre-immunized sera for at least 70 weeks, whereas those in IM-immunized mice were statistically significant until week 9 and no longer statistically significant after that time point (**Fig. 5E**).

Furthermore, we examined the S1-specific IgG1 and IgG2a titers at weeks 5/6, 20, 40, and 70 to determine if there were significant differences due to the route of immunization and longevity. IgG1 and IgG2a EPT of the MAP immunized group were statistically significant compared to pre-immunized controls at some tested time points (**Fig. 5F and 5G**). Interestingly, the mean EPT of IgG1 in MAP ID group increased significantly at later time points, at weeks 40 and 70, while those in the IM group decreased. The mean EPT of IgG2a in the MAP ID group were increased steadily until week 40 and dropped at week 70, while the IM group showed steady mean titers. Based on the ratio of IgG1 to IgG2a antibodies, similar Th2-prevalent responses were observed regardless of vaccine dose and route of immunization at all time points, except for a slight shift towards to a Th2-biased immune response at week 70 in the MAP ID group due to the steady IgG1 and slight drop in IgG2a (**Fig. 5H**). These results suggest that MAP intradermal delivery induces potent and long-lasting SARS-CoV-2 S1-specific antibody responses compared to subunit intramuscular injection, while showing similar Th2-prevalent responses.

### 3.5. Dose sparing and gamma irradiation for sterilization

Next, we investigated the immunogenicity of MAP-rS1RS09 through dose sparing and gamma irradiation (25 kGy) sterilization as part of our efforts toward clinical-grade manufacturing of MAP subunit vaccines. Gamma irradiation has been previously employed as a terminal sterilization method for MAP-SARS-CoV-2 subunit vaccines [37]. For dose sparing, we produced 5, 15, and 45 *μ*g of MAP-rS1RS09 by adjusting the number of microneedle arrays, and some of these MAPs were sterilized by gamma irradiation (**Fig. 6A**). BALB/c mice (5 per group) were primed and boosted at three-week intervals with either non-irradiated (-) or irradiated (+) 5, 15, or 45 *μ*g of MAP-rS1RS09 intradermally (**Fig. 6B**). Serum samples were collected at weeks 0, 3, 6, 9, 12, 16, 28, 52, 71, 90, and 104 after prime immunization, and the S1-specific IgG EPT was examined by ELISA. All vaccinated groups exhibited higher S1-specific IgG EPT at week 6 compared to week 3, with only 15 and 45 *μ*g MAP-rS1RS09 groups showing significant increases. This significant increase was sustained even at week 71 post-prime vaccination when compared to pre-immunized sera, indicating a robust and sustained immunogenic response, consistent with the promising outcomes observed in the earlier experiment (**Fig. 6C**). Interestingly, the MAP-rS1RS09 group maintained a steady geometric mean S1 IgG EPT through week 28, reaching levels as high as the peak at week 6, with minimal waning of the immune response through week 104 post-prime. The mean IgG EPT at the final week 104 showed dose-dependence, although statistical significance was weak due to the low number of animals. Notably, the immunogenicity of gamma irradiation-sterilized MAP vaccines was comparable to that of non-irradiated MAP vaccines in all vaccine groups (**Fig. 6D**). These findings support the feasibility of gamma irradiation as a terminal sterilization approach for our clinical MAP-rS1RS09 vaccines.

**Fig. 6.**
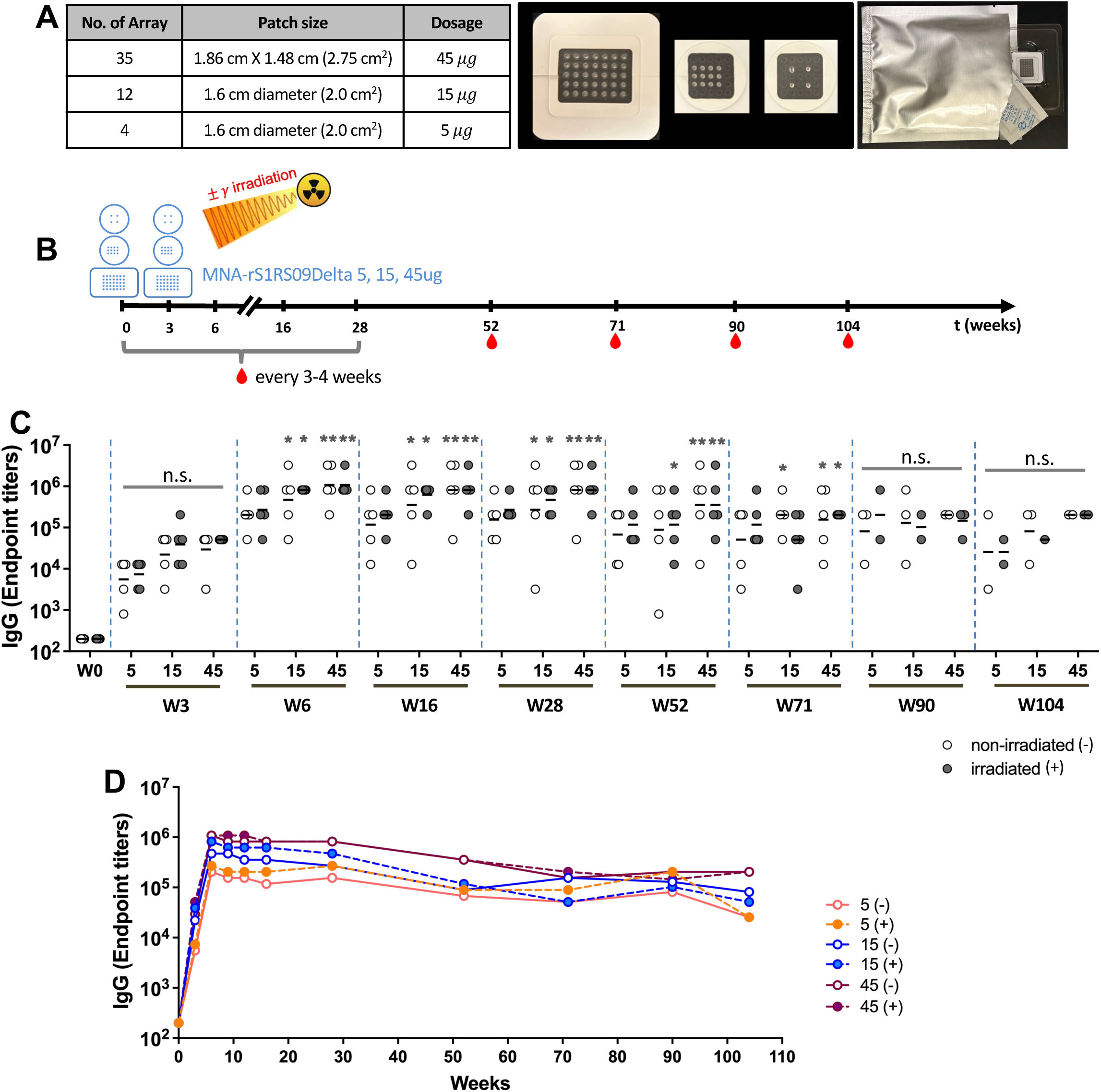
Dose sparing and gamma irradiation responses. (A) Fabrication of dissolving microneedle patch, MAP-rS1RS09 was composed of 35 microneedles. (B) Experimental schedule representing the immunization timeline. Balb/c mice (5 per group) were prime and boosted three weeks intervals with either non-irradiated (-) or irradiated (+) of 5, 15, 45 *μ*g of MAP-rS1RS09Delta. The red drops represented bleeding. (C) Reciprocal serum endpoint dilutions of SARS-CoV-2-S1-specific antibodies were measured by ELISA to determine the IgG endpoint titers at weeks 0, 3, 6, 16, 28, 52, 71, 90, and 104. (D) The SARS-CoV-2-S1-specific IgG titers at all time points until week 104 post-prime (BALB/c (N = 5) except at week 90 (N = 2-3) and at week 104 (N = 1-3)).

We also examined the S1-specific IgG1 and IgG2a titers to determine if there were significant differences in the ratio of these subtypes due to vaccine dose and gamma irradiation. Sera collected at weeks 0, 9, 28, 52, 71, 90, and 104 after prime-boost immunization with either non-irradiated (-) or irradiated (+) of 5 or 45 *μ*g of MAP-rS1RS09 were subjected to isotype-specific ELISA. The IgG1 subclasses of the 5 *μ*g MAP-rS1RS09 group showed a significant increase at weeks 9 in the non-irradiated group and at week 52 in the irradiated group after initial immunization compared to pre-immunized sera (**Fig. 7A**). The IgG2a subclasses of the non-irradiated 5 *μ*g MAP-rS1RS09 group increased significantly at weeks 52 and 71, while those of the irradiated MAP group increased at weeks 28, 52 and 71 (**Fig. 7B**). Interestingly, mean titers of IgG1 and IgG2a in both non-irradiated and irradiated groups remained steady at most time points, with reduced titer at week 104, indicating a shift toward a Th2-biased immune response (**Fig. 7C**).

**Fig. 7.**
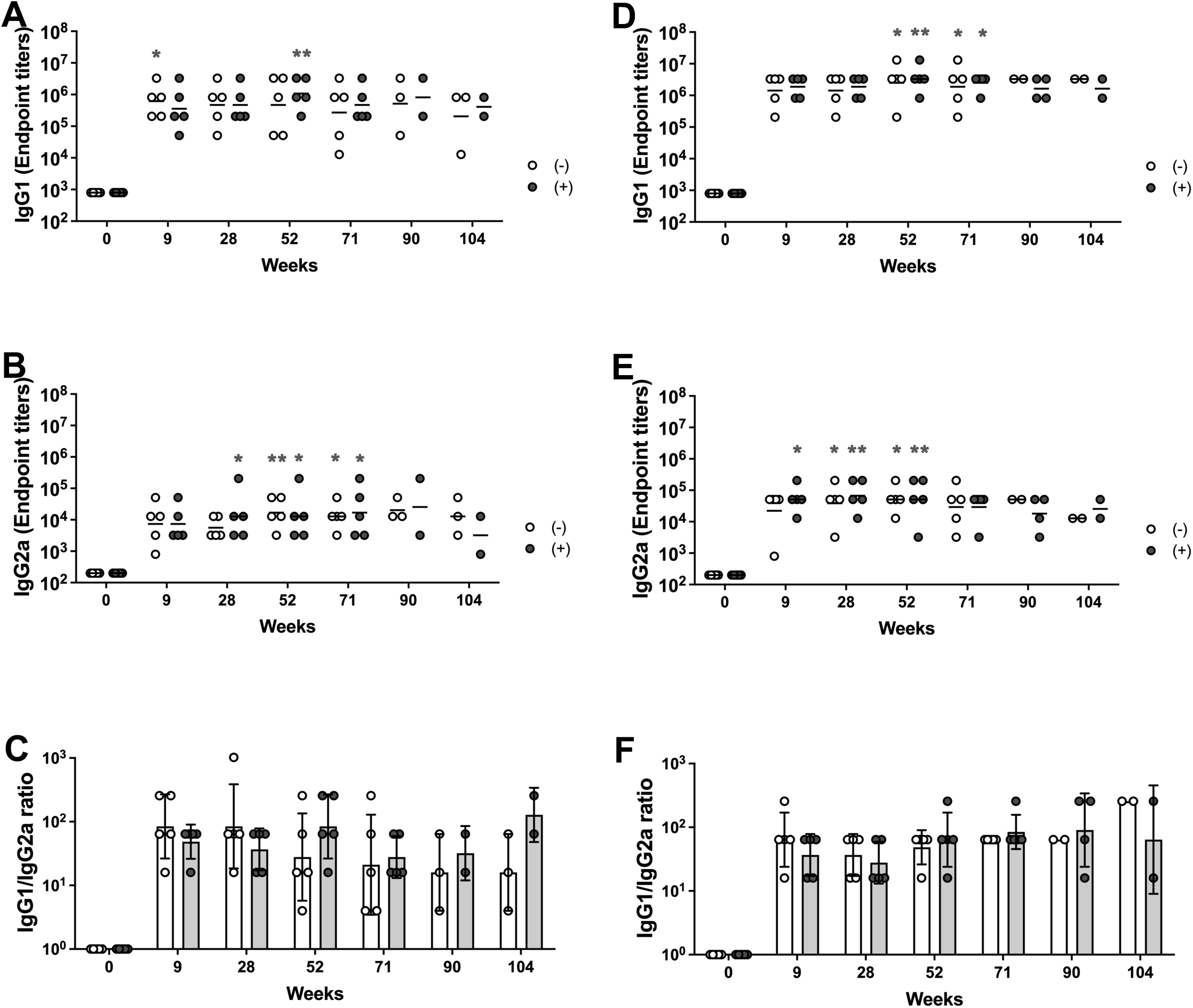
IgG subclasses. Sera were collected from the mice immunized 5 *μ*g of MAP-rS1RS09 with and without irradiation (A, B, and C) or 45ug of MAP-rS1RS09 with and without irradiation (D, E, and F) at weeks 0, 9, 28, 52, 71, 90 and 104. SARS-CoV-2-S1WU-specific IgG1 (A and D) an IgG2a (B and E) were quantified by ELISA to determine each IgG subclasses endpoint titer. The titers at each time points were showed for each mouse. The bars represent geometric mean. (C and F) S1-specific IgG1/IgG2a ratios of individual mice at each time points as mean values with SEM. Groups were compared by the Kruskal-Wallis test at all time points, followed by Dunn’s multiple comparisons. Significant differences are indicated by *p < 0.05, **p < 0.01.

The IgG1 subclasses of both the non-irradiated and irradiated 45 *μ*g MAP-rS1RS09 groups showed a significant increase at weeks 52 and 71 after initial immunization compared to pre-immunized sera (**Fig. 7D)**. The IgG2a subclasses of the non-irradiated 45 *μ*g MAP-rS1RS09 group increased significantly at weeks 28 and 52, while those of the irradiated MAP group increased at weeks 9, 28, and 52 (**Fig. 7E**). Interestingly, the mean titer of IgG1 remained steady at all time points, while that of IgG2a decreased slightly at week 104, indicating a shift toward a Th2-biased immune response (**Fig. 7F**), consistent with the findings in **Fig. 5A and B**. Therefore, a high dose of antigen induced a higher IgG1 titer and a significant IgG2a titer earlier than in the low-dose group. In the case of IgG2a, the irradiated groups induced earlier responses than the non-irradiated groups in both high- and low-dose groups, although there were no significant differences among the groups. Thus, these results suggest that Th2-prevalent responses were observed at all time points based on the ratio of IgG1 to IgG2a antibody subclasses, and there were no significant differences due to vaccine dose or gamma irradiation.

### 3.6. Non-irradiated and irradiated MAP-S1RS09 induce virus neutralizing antibodies

We measured the neutralization capacity of antibodies induced by our vaccine against SARS-CoV-2 Wuhan, Delta, and Omicron variants using a microneutralization assay (NT_90_) (**Fig. 8**). Robust titers of neutralizing antibodies were found against the Wuhan and Delta variant, with the highest levels in the 15 *μ*g and 45 *μ*g dose groups, showing a statistical difference observed only against the Delta variant. Additionally, the 15 *μ*g and 45 *μ*g dose groups were the only ones to show any detectable neutralization activity against the Omicron variant (**Fig. 8**). Notably, the immunogenicity of gamma irradiation-sterilized MAP-S1RS09 vaccines was comparable to that of non-irradiated MAP vaccines in the 15 *μ*g and 45 *μ*g dose groups against the Wuhan and Delta variant, and in the 45 *μ*g dose group against Omicron. However, sera from one to three out of five mice showed neutralization activity in the 5 *μ*g non-irradiated group against the Wuhan and Delta variant, and in the 15 *μ*g irradiated group against Omicron. Altogether, no significant differences were observed between irradiation and non-irradiation in all the groups, thereby supporting the feasibility of gamma irradiation as a terminal sterilization approach for the clinical use of MAP-rS1RS09 vaccines. These findings demonstrate that administering a minimum dose of 15 *μ*g of MAP can generate detectable neutralizing antibodies against various SARS-CoV-2 variants, including Omicron (BA.1).

**Fig.8.**
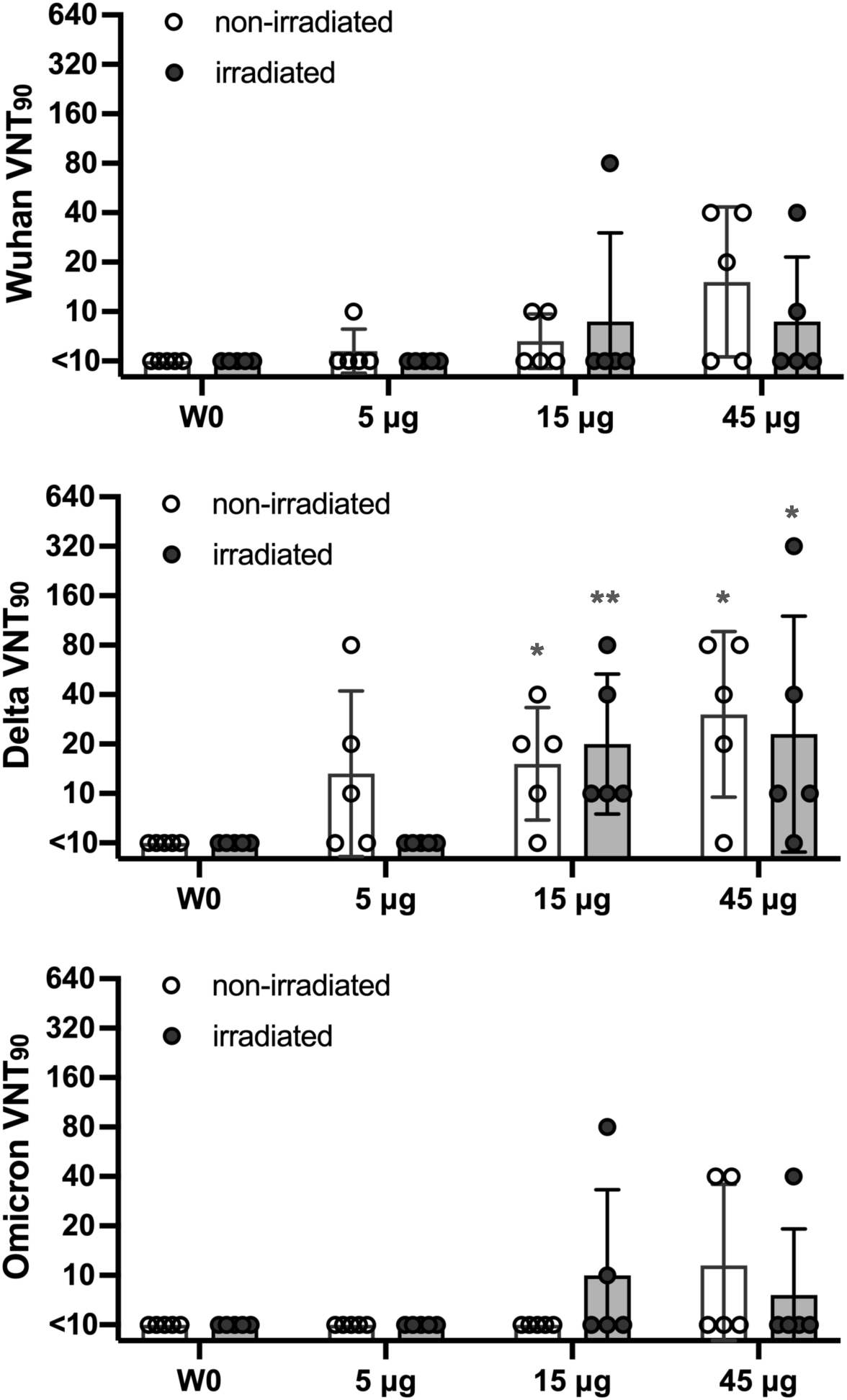
Neutralization of SARS-CoV-2 variants. Serum from mice immunized with SARSCoV-2 S1 via MAP intradermal delivery was assessed using a microneutralization assay (VNT_90_) for neutralization against SARS-CoV-2 variants Wuhan, Delta variant (B.1.617.2), and Omicron variant (BA.1). Serum titers that resulted in 90% reduction in cytopathic effect compared to the virus control were reported. Data are representative of the geometric mean with error bars representing geometric standard deviation. Each group was compared to the pre-immunized sera using the Mann-Whitney test. Significant differences are indicated by **p < 0.01, *p < 0.05.

### 3.7. Long-term Stability of MAP-S1RS09 stored at Elevated Temperature

To assess the stability of irradiated MAP-rS1RS09, patches packed with desiccant were stored either in the refrigerator or at RT for one week, reconstituted in a buffer by shaking for 30 minutes, and compared for protein degradation (**Fig 9A**). rS1RS09 from the irradiated patches showed slight smearing in both temperature conditions in the silver-stained gel after SDS-PAGE (**Fig 9B**). To further assess the effects of extended storage time, we stored the patches for nineteen months at either refrigerator or RT, and investigated protein degradation by measuring the relative smear ratio compared to recombinant proteins stored at −20°C for 19 months in parallel. As shown in **Fig. 9C**, there are no differences of rS1RS09 in MAP stored for nineteen months in both temperature conditions compared to rS1RS09 in MAP stored for one week (**Fig 9B**) or rS1RS09 stored at − 20 ° C for nineteen months, suggesting excellent stability (**Supplementary Fig. 1A**). The mean percentage of protein degradation based on relative intensity (RI) of non-irradiated and irradiated patches using ImageJ analysis was 0.3% and 1.95% when stored at 4°C for nineteen months, and 1.85% and 3.25% when stored at RT, respectively (**Fig 9D**). The degree of protein degradation in irradiated MAPs was comparable to that in non-irradiated MAPs, with no significant differences (1.40% vs. 1.55%), indicating notable stability at RT as good as at 4°C. Furthermore, to assess the stability of long-term storage, the packed MAPs stored at 4°C for nineteen months were further stored under three different conditions: at 4 °C in a refrigerator, at RT on a lab bench, and at 42 °C in a temperature-controlled incubator for one month (**Fig. 9A**). There were no significant differences of rS1RS09 in MAPs depending on temperature conditions compared to non-irradiated MAPs stored at 4°C for nineteen months (**Fig. 9E and Supplementary Fig. 1B**). The percentages of protein degradation in both non-irradiated and irradiated MAPs were on average 0.64% ± 0.64 at 4°C, 1.60% ± 0.39 at RT, and 2.62% ± 0.03 at 42°C, respectively. In contrast, storage of rS1RS09 in buffer solution at 42 °C for one month led to loss of 24.28±0.66% of proteins, whereas rS1RS09 in MAP showed a loss of 2.80±0.15%, based on silver-staining followed by ImageJ analysis (**Fig. 9F and 9G**). The findings revealed that the MAP platform storage of rS1RS09 is stable for at least twenty months without refrigeration, indicating the temperature stability of our gamma irradiation sterilized MAP-rS1RS09 vaccines without dependence on the cold chain.

**Fig. 9.**
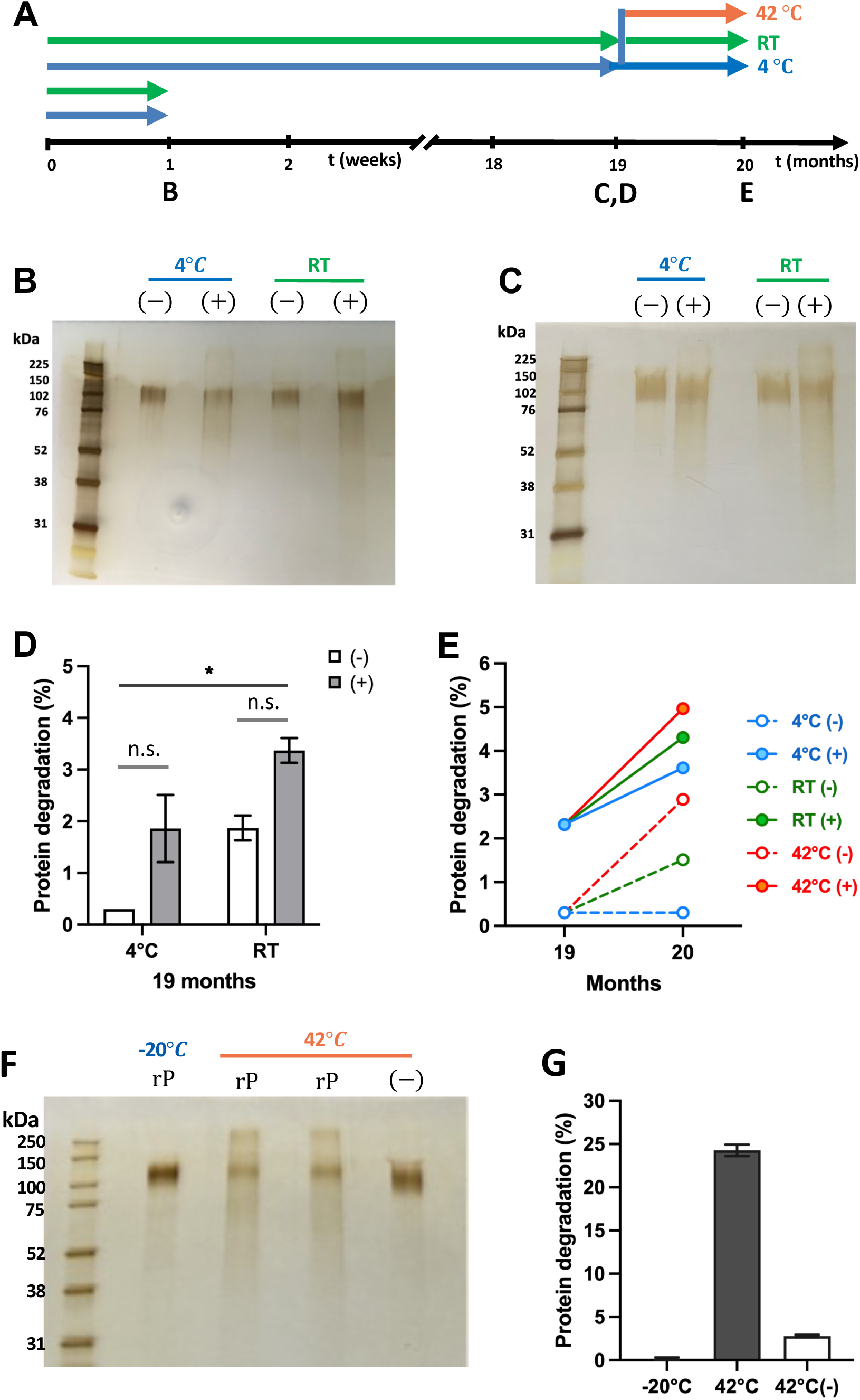
Stability persistence of irradiated MAP. (A) Experimental schedule representing storage temperature and the reconstruction timeline of MAP. Silver staining of the recombinant proteins reconstructed from the non-irradiated (–) and irradiated (+) MAP stored at either 4°C or RT for 1 week (B) and 19 months (C). (D) Percentage of protein degradation in the MAP from silver staining in Fig. 9C, measured by ImageJ. The recombinant protein stored at – 20°C for 19 months was set to 0%. (E) Percentage of protein degradation for an additional one month at different temperatures with the MAP stored for 19 months at 4°C, measured by ImageJ. The recombinant protein stored at –20°C for 19 months was set to 0%. (F) Silver staining of the recombinant proteins (rP) stored at –20°C for 20 months (lane 1), in PBS (lane 2) or in the preservative buffer (lane 3) stored at 42°C for 1 month after 19 months at –20°C, and rP reconstituted from the non-irradiated (–) MAP stored at 42°C for 1 month after 19 months at 4°C (lane 4). (G) Percentage of protein degradation of rS1RS09 in the buffer solution or MAP from silver staining in Fig. 9F, measured by ImageJ.

## 4. Discussion

In our prior research, we established that a SARS-CoV-2 S1 subunit MAP vaccine effectively triggered the S1-specific immune system in mice [37] and the second and third boosts elicited robust antibody responses against S1, effectively neutralizing the virus, resulting in enduring high-level, long-lasting antibody responses [25].

In this study, we found that the SARS-CoV-2 S1 monomer in the presence of RS09 (rS1RS09) is more immunogenic than the trimeric S1 subunit protein, regardless of RS09, considering it as an ideal vaccine candidate. rS1RS09 induced a high level of IgG titer, capable of binding spike proteins expressed on the cell surface, mimicking natural viral trimeric spike proteins. Furthermore, rS1RS09 induced not only a greater IgG titer but also neutralizing antibodies, as determined by the ACE2 binding inhibition assay (**Fig. 1-3**). RS09, TLR4 agonist peptide motif, was identified as an adjuvant using phage display combinatorial peptide technology, which functionally mimicked lipid polysaccharide (LPS), activated NF-κ B signaling, and induced inflammatory cytokine secretion [38]. Interestingly, low (0.1 and 1 *μg*) and high dose (10 and 100 *μg*) of inhaled LPS in a mouse model of asthma induced Th2 and Th1 responses to allergens, respectively [39,40]. Low-dose LPS induced TNF-*α* secretion, which is a key upstream molecule of Th2 polarization, while high-dose LPS induced IFN-*γ*, which is a key role in up-regulating allergen-specific IgG2a, thus restoring Th1 response. Similarly, our present study found that rS1RS09 induced a more toward to Th2-biased immune response compared S1 alone, especially at week 18 post-prime, as evidenced by the increase in IgG1 level (**Fig.2C and 2E**). This phenomenon might be attributed to the low dose of RS09, approximately 0.39 *μg* RS09 from the 45 *μg* of rS1RS09. In a past study study, RS09 reported firstly as an adjuvant was injected at 25 *μg* [38], and 10 *μg* of RS09 adjuvant with a tetravalent DENV nanoparticle vaccine stimulated high IgG2a titers in BALB/c mice [41]. Likewise, antigen containing triple RS09 motif at the C-terminus can impart TLR4 agonist activity, which is approximately 10.85 *μg* from the 100 *μg* of the artificial antigen [42]. This suggests that further evaluation of the molecular context of the TLR4 agonistic motif in rS1RS09 should be required using HEK-Blue cells, derived from HEK293 cells carrying human TLR4/MD2/CD14 gene and a SEAP (secreted embryonic alkaline phosphatase gene) reporter construct inducible by NF-κB signaling [42].

Furthermore, the present data showed that the monomeric form of S1 induced a higher IgG titer compared to the trimeric form regardless of the RS09 motif (**Fig. 2B and 2F**). This might be explained by many epitopes being hidden in the trimeric form compared to the monomeric form, similar to HIV-1, where its most conserved epitopes are concealed inside the Env core and are only exposed after CD4 receptor engagement [43]. Indeed, the epitope sites are not hidden in the isolated RBD construct, but in the Spike trimer, and the RBDs must move from a down- to an up- position, and antibody binding is further hindered spatially by the N-terminal domain (NTD) and the S2 subunit [44].

We also compared the antibody responses induced by MAP of the S1 subunit antigen immunization to those elicited by conventional intramuscular (IM) injections. Despite of no significant difference between the two groups with a low dose of 5 *μ*g rS1(Wu+Beta), the delivery of protein subunit vaccine though MAP is superior to conventional intramuscular injection, although the actual amount of delivered antigen is lower, with an average of 70% [25] (**Fig.4**). Vaccine-induced protection against SARS-CoV-2 diminishes over time, leading to instances of SARS-CoV-2 reinfection [45–47]. Consequently, the challenge of sustaining long-term immunity is the major obstacles in enhancing and approving COVID-19 vaccines [48]. While it’s complex to directly compare human and mouse lifespans, the enduring immune responses observed in this study following additional MAP boosts suggest a potential solution to overcome these hurdles. Interestingly, all groups showed similar Th2-prevalent responses regardless vaccine dose and route of immunization (**Fig. 4F**). In contrast, with a high dose of 45 *μ*g rS1RS09, all vaccinated groups had significantly high and similar geometric mean S1 IgG endpoint titer (EPT) after boost when compared to week 0. The IgG geometric mean of the IM group peaked at week 5 and waned the immune response through week 71, while that of the MAP-rS1RS09 group reached a steady geometric mean S1 IgG EPT through week 71 as high as the peak at week 6, showing a significant difference between two groups (**Fig. 5**). The IgG1 and IgG2a EPT of the MAP group were statistically significant compared to pre-immunized controls and showed similar Th2-baised immune response except later time point, at week 70, leaning more toward to Th2-baised in the MAP group. These results are in line with those of an inactivated influenza vaccine encapsulated in MAP, which induced a much higher IgG1 titer and lower IgG2a titer compared to the intramuscular injection of a liquid subunit vaccine [49].

Nevertheless, our research revealed that rS1RS09 encapsulated in MAP induced a superior immune response compared to conventional IM injection (**Fig. 4 and 5**). A previous study had shown that gamma irradiation has been employed as a terminal sterilization method for MAP-SARS-CoV-2 subunit vaccines [37]. As part of our efforts toward clinical-grade manufacturing of MAP subunit vaccines, further investigation of dose sparing and gamma irradiation (25 kGy) sterilization was performed. As shown in Fig 6, 15 and 45 *μ*g of MAP-rS1RS09 groups induced and sustained significant IgG titer even at week 71 post-prime vaccination when compared to pre-immunized sera. The immunogenicity of gamma irradiation sterilized MAP vaccines was comparable to that of non-irradiated MAP vaccines in all vaccine groups (**Fig. 6C**). Thereby, these findings support the feasibility of gamma irradiation as a terminal sterilization approach for the clinical usage of our MAP-rS1RS09 vaccines.

We also examined the S1-specific IgG1 and IgG2a titers to determine if there were significant differences in the ratio of these subtypes due to vaccine dose and gamma irradiation by comparison with 45 *μ*g and 5 *μ*g groups. Critically, a 45 *μ*g dose of antigen induced a higher IgG1 titer and a significant IgG2a titer earlier than in a 5 *μ*g dose group, resulting in a forward

Th2-biased response (**Fig.7**). Furthermore, the irradiated groups induced earlier responses than non-irradiated groups in both high and low dose groups, although there were no significant differences among the groups, consistent with previous reports that Malaria SPf66 synthetic peptide release rate from poly (lactic-co-glycolic acid) (PLGA) microspheres was slightly faster after gamma-irradiation, although no apparent effect on SPf66 integrity and formulation properties, such as morphology, size, and loading [50]. Subcutaneous administration of irradiated and non-irradiated microspheres into mice induced a similar immune response (IgG, IgG1, IgG2a levels) [50]

Notably, gamma irradiation-sterilized MAP vaccines induced comparable neutralizing antibodies against three SARS-CoV-2 variants to that of non-irradiated MAP vaccines, with no statistical significance among groups, conforming that the immunogenic activity of the antigen is maintained after gamma-irradiation (**Fig. 8**). Critically, sera from one and three out of 5 mice showed neutralization activity in the non-irradiated 5 *μ*g group against the Wuhan and Delta variants, respectively, whereas the irradiated 5 *μ*g group was under the limit of detection. In contrast, sera from two out of 5 mice showed neutralization activity in the irradiated 15 *μ*g group against Omicron variant, whereas the non-irradiated 15 *μ*g group was under the limit of detection, indicating few differences in antigenicity by gamma-irradiation. Several previous studies have reported that irradiation would cause breakage and polymerization of peptide chains, and glycosylation modification of proteins [51,52]. Irradiation treatment effects on the protein structure and digestion characteristics of seed-watermelon (Citrullus lanatus var.) kernel protein depending on doses of irradiation. After the irradiation treatment, the kernel protein was unfolded by decrease in alpha-helix and beta-sheet structure, and an increase in coil structure, resulting in the exposure of hydrophobic groups [53]. Moreover, gamma irradiation induces protein changes that enhance immunogenicity for snake venoms, *Coccidian* parasites, and *Toxoplasma gondii* protein extracts [54,55]. Therefore, these findings support that the little change in antigenicity by gamma irradiation as a terminal sterilization approach may be beneficial for neutralizing a broad spectrum of emerging variants. Under detectable neutralizing activity in low-dose MAP vaccine can be overcome through additional boost or combinations with adequate adjuvants such as Poly(I:C), monophosphoryl lipid A, cholera toxin, zymosan [15,56–58].

We also demonstrated that the MAP platform storage of rS1RS09 is stable for at least twenty months without refrigeration, indicating the temperature stability of our gamma irradiation-sterilized MAP-rS1RS09 vaccines without dependence on the cold chain (**Fig. 9**). Furthermore, an additional one-month storage at high temperature showed that superior benefits of the MAP platform compared to the solution (**Fig. 9F and 9G**). Similarly, recent studies have shown that MAP for administration of inactivated influenza or inactivated polio vaccine improved thermal stability compared with conventional liquid vaccines, which could enable the distribution of vaccines with less reliance on cold chain storage [49,59]. Moreover, researchers found that hyaluronic acid (HA) plays an antioxidant role not only in patches but also in the solution during the manufacturing process and prevents reductions in AA2G content by irradiation [60]. Furthering this, gamma irradiation affected the mechanical properties and architecture of the needles of carboxymethyl cellulose (CMC), but not HA-based patch [61]. They also suggested that e-beam (40 kGy), which provides lower penetration and higher doses into sterilizing materials, is suitable for the terminal sterilization of HA-based DMN patches, because gamma-ray (20 and 30 kGy) irradiation significantly degraded the encapsulated AA2G, while e-beam maintained AA2G activity [60]. However, AA2G is a vitamin C derivative, which is expected to be easily oxidized by the radicals generated from irradiation. From the perspective of immunogenicity, gamma irradiation of antigens can mimic the promotion of slow release by generating smaller solubility protein aggregates like an adjuvant or oxidizing the antigens, as similarly reported for neutrophil myeloperoxidase in acute inflammation [62–64]. Gamma irradiation-inactivated (RI) Respiratory Syncytial Virus (RSV) vaccine exacerbates pulmonary inflammation by switching from prefusion to postfusion F protein. RI-RSV caused more severe ERD than did formalin-inactivated (FI)-RSV, and these immune responses are likely due to pre-F to post-F conformational changes by radiation-induced redox oxygen species (ROS) [65]. Moreover, gamma irradiation of antigens also induced a better cellular immune response [66]. In addition, irradiated proteins undergo one of the chemical changes, such as cross-linking, which can support the chemical bonding of HA to antigens, stimulating strong and enduring humoral responses like an adjuvant [67]. Indeed, in this study, the GMT of IgG in all irradiated groups was observed to be slightly higher than those induced in the non-irradiated groups at most time points (**Fig.6B**).

This study had two limitations, which will be addressed in future research. These include conducting a SARS-CoV-2 challenge study to evaluate the protective effectiveness of non-irradiated and irradiated MAP vaccination and assessing the immunogenicity of non-irradiated and irradiated MAP-S1RS09Delta stored for the long-term at different temperatures compared to those stored for the short-term. Interestingly, a study has reported that IgG titers were similar in mice vaccinated with MAP encapsuled inactivated influenza vaccine stored for one month at 4℃, 25℃, and 45℃ compared to MAP vaccine stored for one day at 4℃, leading to complete protection against the virus challenge with no significant weight loss [49].

In summary, our research revealed that delivery with MAP platforms for subunit vaccine is superior to conventional IM injection in terms of immunogenicity and long-term stability. We investigated the impact of gamma irradiation as a terminal sterilization method for MAP platforms targeting SARS-CoV-2 S1 antigens, resulting in the development of comparable antibody responses. These findings suggest the potential for further research into utilizing irradiated MAP to deliver S1 subunit vaccines for emerging SARS-CoV-2 variants. Such vaccines could serve as safe boosters to enhance cross-neutralizing antibody responses. This approach holds promise as a viable platform for facilitating widespread global vaccination efforts.

## 5. Conclusions

This study presents the first assessment of long-term immunogenicity and stability of gamma-irradiation-sterilized dissolving microarray patches (MAP) for the SARS-CoV-2 subunit vaccine. rS1RS09 was identified as an ideal immunogen after investigating the immunogenicity of four recombinant Delta S1 proteins (rS1, rS1RS09, rS1f, and rS1fRS09) in monomeric or trimeric forms in BALB/c mice. Moreover, MAP delivery of rS1RS09 elicited strong and sustained antibody responses, outperforming conventional intramuscular injections. Both irradiated and non-irradiated MAP vaccines showed comparable immunogenicity, demonstrating a robust, long-lasting immunity and a dose-sparing effect. The enhanced stability of the subunit vaccine in MAPs during extended storage outside the cold chain supports their potential for global distribution and holds promise as an effective, stable, and scalable SARS-CoV-2 vaccine candidate. Overall, gamma irradiation as a terminal sterilization method is feasible for mass production and future pandemic preparedness.

