## Supplemental Fig 1 for "Long-term Immunity of a Microneedle Array Patch of SARS-CoV-2 S1 Protein Subunit Vaccine Irradiated by Gamma Rays in Mice"

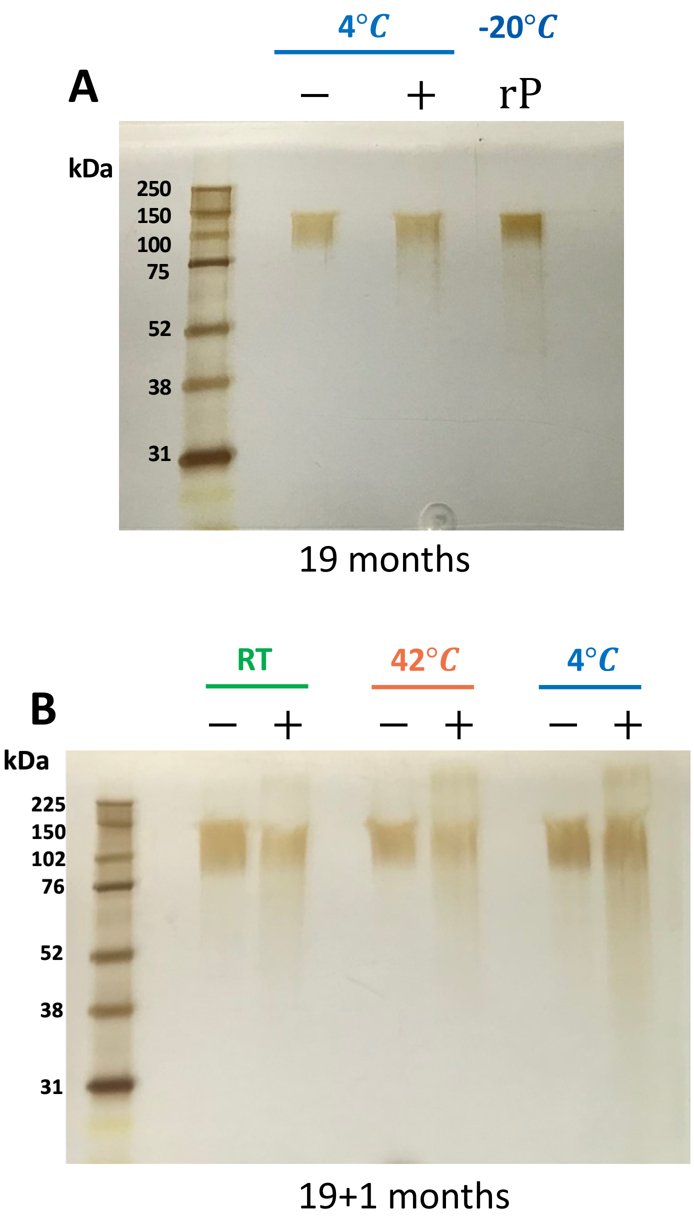


**Supplementary Fig. 1. Stability of Irradiated MAP over time** (A) Silver staining of the recombinant proteins (rP) reconstructed from the non-irradiated (–) and irradiated (+) MAP stored at 4°C for 19 months and recombinant protein (rP) stored at -20°$C$ for 19 months (B) rS1RS09 reconstructed from the non-irradiated (–) and irradiated (+) MAP stored for an additional one month at 4°C, RT, and 42°C, after 19 months at 4°C. Approximately 200ng (A) and 300ng (B) of rP were loaded.
